# scStudio: A User-Friendly Web Application Empowering Non-Computational Users with Intuitive scRNA-seq Data Analysis

**DOI:** 10.1101/2025.04.17.649161

**Authors:** Marta Bica, Karine Serre, Nuno L. Barbosa-Morais

**Author notes:** Corresponding authors: Marta Bica, Nuno Barbosa-Morais.

## Abstract

**Background:** Single-cell RNA sequencing (scRNA-seq) has revolutionized our understanding of cellular heterogeneity by providing detailed insights into gene expression at the individual cell level. Despite its potential, the complexity of scRNA-seq data analysis often poses challenges for researchers without computational expertise.

**Findings:** To address this, we developed scStudio, a user-friendly, comprehensive, and modular web-based application designed to democratize scRNA-seq data analysis. scStudio is equipped with a suite of features designed to streamline data retrieval and analysis with both flexibility and ease, including automated dataset retrieval from the Gene Expression Omnibus. Users can also upload their own datasets in a variety of formats, integrate multiple datasets, and tailor their analyses using a wide range of flexible methods with options for parameter optimization. The application supports all the essential steps required for scRNA-seq data analysis, including in-depth quality control, normalization, dimensionality reduction, clustering, differential expression, and functional enrichment analysis. scStudio also tracks the history of analyses, supports session data storage and export, and facilitates collaboration through data sharing features.

**Conclusion:** By developing scStudio as a user-friendly interface and scalable architecture, we address the evolving needs of scRNA-seq research, making advanced data analysis accessible and manageable while accommodating future developments in the field. scStudio is freely available at https://compbio.imm.medicina.ulisboa.pt/app/scStudio.

## Introduction

Single-cell RNA sequencing (scRNA-seq) is a powerful technology that enables the analysis of gene expression at the individual cell level [1]. Unlike traditional RNA sequencing, which measures the average gene expression of a population of cells and obscures cellular diversity, scRNA-seq provides a detailed view of the distribution of gene expression levels across a population of individual cells, enabling the exploration of complex and heterogeneous tissues [1]. This technique has revolutionized our understanding of biological processes by uncovering diverse cell types [1,2], identifying rare cell populations [1,3], and capturing dynamic gene expression changes during development [1,4], disease progression [1,4,5], and response to stimuli [1]. By dissecting the transcriptomic landscape at the single-cell level, researchers can explore cellular heterogeneity, revealing key regulatory mechanisms and therapeutic targets with potential far-reaching implications for both basic and clinical research. The utilization of scRNA-seq protocols has hence been rapidly expanding across various research fields, including developmental biology, immunology, neuroscience, cancer, and aging research [1]. The generation of scRNA-seq data is experiencing significant growth, a trend that shows no signs of slowing down [6,7]. Since 2013, the publication rate of scRNA-seq experiments has surged, from one study per month to over 50 studies per month by early 2020 [7]. More recently, journals have begun requiring that generated datasets be published in public repositories such as NCBI’s Gene Expression Omnibus (GEO) [8]. This mandate is crucial for enabling the scientific community to scrutinize results through replication and validation, as well as for the potential application of new discoveries. Many scRNA-seq datasets remain underexplored, presenting opportunities for reuse in addressing other biological questions [9]. Despite the accumulation of scRNA-seq datasets in public repositories like GEO [8], most single-cell data analysis tools still require programming knowledge, typically in R or Python, hindering their utilization by scientists without computational expertise.

To address this barrier, several applications with visual interfaces for scRNA-seq data analysis have been developed (Supp. Table 1). However, these tools often present at least one of the following shortfalls, preventing them from fully meeting the needs of non-expert users:

1. **Limited selection of datasets for exploration**: programs including only a restricted group of datasets pre-processed by the developers, or a small manually curated database that rapidly gets outdated, given the accumulation of newly generated data.
2. **No automated access to publicly available data**: instead of automatically retrieving data from public repositories, users need to upload to the program a count matrix and cell-level annotations; obtaining these from repositories is cumbersome because there is no standard file format or naming convention for deposited data.
3. **Lack of flexibility**: data are pre-processed using parameters decided based on prior analyses by the developers (e.g., fixed clustering resolution, differential expression analysis between fixed groups of cells), restricting data exploration’s scope.
4. **Limited diversity of data exploration methods**, often dependent on deprecated software needing frequent updates.
5. No support for multiple dataset integration and meta-analyses.
6. **Requirement of local software installation,** often unfeasible due to some computational resource-demanding and thereby expensive tools.
7. **“Black boxes”, paid and/or closed-source software**: programs with restricted availability and little transparency of algorithms and parameters employed.

We have thus developed scStudio, a flexible user-friendly web-based application that goes beyond visualization of fixed pre-processed data and empowers non-computational scientists to perform sound and intelligible analyses of own and publicly available scRNA-seq datasets. Our tool allows performing interactive analyses, including in-depth quality control (QC), dimensionality reduction, clustering, differential gene expression and functional enrichment analysis. scStudio was built upon four main pillars, namely:

### Creating a portal for GEO single-cell data analysis

GEO is a public repository that archives and freely distributes comprehensive sets of next-generation sequencing, and other forms of high-throughput functional genomic data submitted by the scientific community [8]. GEO is the most used repository for scRNA-seq data, storing 78.1% of single-cell studies in curated public databases [10]. scStudio users are able to directly access GEO data through an interface performing programmatic calls with user-selected GEO IDs (e.g., GSE137710) to automatically upload read count matrices and available metadata, assuming that datasets follow minimal format compliance rules, discussed further in this article.

### Implementing a decision support system, avoiding “black boxes”

scStudio has an interface to run scRNA-seq data analysis pipelines in a tutorial-like conceptual navigation of each step. Automated pre-processing and QC of single-cell data are achievable with algorithms designed for specific protocols (e.g., doublet identification for droplet data) but our goal is to provide users with high-quality visualization and analytical tools to support their decision-making at each analysis stage. One example is the construction of clustering trees [11] to inspect the variation of clustering results across different resolutions, alongside with marker genes for each cluster, and thereby fine-tune the definition of clusters of cells that, at different resolutions, likely represent true biological variability. Another example is the inclusion of methods for functional enrichment analysis [12] and visualization of results.

### Data integration and sharing

scStudio’s flexibility expands both the resolution of analyses (e.g., by allowing users to select subsets of the dataset and re-running the pipeline) and their scale, by enabling, with robust integration algorithms [13], joint analysis of multiple datasets (both public and own) for exploration of cell types and states across studies, ultimately contributing to novel cell atlases. Finally, users are able to save their work sessions at any analysis stage, avoiding re-running computationally expensive jobs, and directly share annotated datasets, facilitating reproducibility and scrutiny of analyses by fellow scientists.

### Increasing diversity and flexibility of scRNA-seq analysis methods

As the field of scRNA-seq continues to advance, the range of available analysis methods has grown substantially [13], reflecting the diverse goals of the research community. These tools have enabled researchers to explore cellular heterogeneity, gene regulation, and dynamic biological processes. As shown by Svensson *et al.* [7], by tracking the types of analyses performed with scRNA-seq data, we can gain insights into the community’s broader research objectives. Clustering cells based on gene expression to identify molecular cell types remains the most common application, featured in nearly 90% of studies [6, 7]. Visualization techniques, particularly t-distributed Stochastic Neighbor Embedding (t-SNE) [14], became widely adopted for single-cell analysis since its introduction in 2015 [7]. However, its usage has slightly declined recently with the emergence of the Uniform Manifold Approximation and Projection (UMAP) method [15,7]. Another popular method is pseudotime analysis, used in about half of all studies to investigate developmental trajectories and dynamic processes over time [7]. Despite the growing diversity of these methods, their complexity often presents challenges, particularly for non-computational scientists. On one hand, traditional analysis pipelines require a solid understanding of programming languages, creating barriers to accessibility. On the other, those methods’ underlying assumptions and theoretical concepts are not intelligible to most users, making the results of their application prone to misinterpretation [16]. To address these issues, we developed scStudio not only to support a wide range of scRNA-seq analysis methods but also to offer a user-friendly interface that empowers researchers from various backgrounds to perform advanced analyses with minimal computational expertise. By integrating these diverse and flexible analysis options, scStudio serves as a comprehensive platform that bridges the gap between cutting-edge bioinformatics and accessible data analysis, making the power of scRNA-seq available to a broader scientific audience.

## Methods

### Development and framework

scStudio was developed as a web-based application using the R Shiny package [17] (version 1.10.0), that provides a platform for building interactive web interfaces.

### Modular architecture

We designed scStudio grounded in a modular architecture that emphasizes both flexibility and scalability. This modularity is reflected in several key aspects of the application, allowing for the seamless integration of new analytical methods and ensuring that users can easily manage and save their data across different stages of analysis.

### Multi-app functionality and method-specific tabs

Each main analysis step is implemented as a separate Shiny app, with its own app.R file. This modular approach allows users to run multiple types of analyses simultaneously, each in its own dedicated app browser window. This design improves overall efficiency and user experience by enabling concurrent analysis tasks. For each main step, scStudio is organized around a tabbed interface, with each tab dedicated to a specific method, e.g., Gene Set Enrichment Analysis (GSEA) [12] in the functional enrichment analysis (FEA) step. For each method, a separate User Interface (UI) and server file has been created, encapsulating the specific functions and logic required to run the analysis. This approach isolates the functionality of each method, ensuring that adding new methods does not interfere with existing ones. This modular separation also simplifies the process of updating or modifying individual methods without affecting the overall structure of the application.

Beyond their modular design, the applications remain interconnected through periodic md5sum-based integrity checks to detect modifications in session-relevant objects. This mechanism allows updated data to be incorporated dynamically without disrupting the current session state. For instance, the generation of a new normalized count matrix or the completion of a t-SNE analysis with alternative parameters in one module is automatically detected and made available as a selectable input in other modules once the background process has completed.

### Data storage and management

In addition to the modular approach to UI and server functions, we employed a modular data storage system in scStudio. Data generated by different analytical methods are stored in separate .rds files, corresponding to the specific type of analysis. This compartmentalized storage strategy not only ensures that data are organized and easy to access but also supports the flexibility to accommodate new data formats as additional methods are integrated into the app. This modular data management system is particularly advantageous in a research environment where multiple types of analyses are performed on the same dataset. By storing results separately for each method, scStudio allows users to save and revisit their analyses without the risk of data corruption or loss from overwriting files. Additionally, as new methods are developed, they can be easily incorporated into the existing framework by simply adding a new .rds file, without disrupting the current workflow or data structure.

### Gene Expression Omnibus interface

We used the GEOquery package [19] (version 2.62.2) to facilitate automatic retrieval of GEO data based on user-selected GEO IDs [8]. Specifically, the getGEOSuppFiles function was employed to download supplementary files typically associated with GEO entries for scRNA-seq datasets. To accommodate the diverse file formats commonly encountered in GEO for scRNA-seq data entries, we developed three distinct workflows:

- Workflow 1 for tabular data formats, including comma-separated values (CSV), tab-separated values (TSV), plain text (TXT), and Excel files (XLSX).
- Workflow 2 for Cell Ranger Market Exchange Format (MEX) files, often used to store sparse matrices in scRNA-seq data generated with 10X Genomics technology [20].
- Workflow 3 for HDF5 (.h5) files, commonly used for storing large datasets efficiently.

We designed this system with flexibility, making it easy to incorporate further workflows to accommodate other data types as needed.

### Quality control

Poor quality cells can be filtered based on user-provided parameters, namely, library size, number of unique features detected, and percentage of reads aligned to mitochondrial genes. Users can compute 𝑙𝑜𝑔_2_ or 𝑙𝑜𝑔_10_ transformed metrics to assess deviations from the normal distribution. Cells can either be immediately removed from the dataset or flagged for downstream inspection. Users are also able to inspect all cells or segregate them by variables available in the dataset’s metadata e.g., to detect poor-quality samples. Gene filtering is also available, allowing filtering of genes based on a user-provided minimum number of total counts and number of expressing cells. Furthermore, we used the scDblFinder package [21] (version 1.8.0) to allow the identification of doublets, which are typically present in droplet-based data.

### Normalization and batch effect correction

We currently provide two distinct methods for library size (sequencing depth) normalization, scran [22] (version 1.22.1) and SCTransform [23] (version 0.4.1). In the future, we plan to expand this list to include additional normalization methods that may perform better with specific protocols or dataset designs. For batch effect correction, we implemented the removeBatchEffect function from the limma package [24] (version 3.50.3). To evaluate the proportion of variance in the count matrix that can be attributed to each known variable in the dataset, we implemented a variance explained plot using the getVarianceExplained function from the scater package [25] (version 1.22.0).

### Feature selection and dimensionality reduction

We used the modelGeneVar function from the scran package [22] (version 1.22.1) to identify highly variable genes (HVGs) using a modeled mean-variance trend [22]. The top HVGs can then be employed in downstream analyses, including dimensionality reduction and clustering, to mitigate noise from lowly expressed genes and/or genes with minimal biological variance.

For Principal Component Analysis (PCA) [26], we used the runPCA function from the BioSingular package [27] (version 1.10.0), with automatic centering of the data and scaling as an optional step. For t-SNE [14], we employed the calculateTSNE function from the scater package [25] (version 1.22.0). The perplexity parameter can be adjusted within a range that allows users to explore its impact, with low perplexity emphasizing the preservation of local structures and high perplexity focusing on global structure [14]. Additionally, we implemented UMAP [15] using the calculateUMAP function from the scater package [25] (version 1.22.0). The parameters minimum distance and number of neighbors that control emphasis on local versus global structure [15] are both adjustable, enabling users to explore, compare, and optimize them based on their specific data and biological questions (e.g., the number of expected cell types or a focus on broader cell populations *versus* more subtle cell states). These dimensionality reduction methods can be applied to raw and/or corrected count matrices to evaluate the effects of normalization and batch effect correction.

### Clustering analysis

We used the FindNeighbors and FindClusters functions from the Seurat package [28] (version 4.1.1) to allow users to obtain cell clusters across a range of resolutions from 0.1 to 2, incremented by 0.1. Briefly, Seurat employs a graph-based clustering approach where a shared nearest neighbor (SNN) graph is constructed based on the chosen feature space [28]. Edges in the graph represent similarities between cells, and clusters are identified using the Louvain algorithm to optimize modularity and define communities of highly similar cells [28]. Users can choose the space in which clustering is performed, either by directly applying it to the count matrix with a selected set of features (e.g., HVGs) or by using a precomputed PCA representation with a user-defined number of principal components. A clustering tree is generated using the clustree function from the clustree package [11] (version 0.5.1). Users can then identify marker genes for their chosen clustering resolutions by performing differential expression analysis, comparing each cluster against the others. This analysis is performed using the FindAllMarkers function from the Seurat package [28] (version 4.1.1). Users can set a minimum absolute average fold-difference (log-scale) between groups as a gene filtering criterion e.g., to increase the speed of the computation. Additionally, multiple statistical tests can be selected to benchmark differences between methods, including the Wilcoxon Rank Sum test, Likelihood-ratio test, Receiver Operating Characteristic analysis (ROC), Student’s t-test, and a hurdle model specifically designed for scRNA-seq data (MAST) [29]. The final output includes the following metrics:

- Adjusted p-value using the Bonferroni method.
- Percentage of cells that express the gene in the cluster (Target) and in the remaining cells (Other).
- Average fold-difference in log scale, computed using either the mean or median gene expression for each group, 𝑙𝑜𝑔_2_𝐹𝐶(𝑚𝑒𝑎𝑛) or 𝑙𝑜𝑔_2_𝐹𝐶(𝑚𝑒𝑑𝑖𝑎𝑛), respectively. We included median-based log_2_FC as a complementary metric due to its robustness to outliers and dropout events, common features of skewed and zero-inflated distributions, thereby better capturing consistent expression shifts across the target cell population. Briefly, for each gene, we calculate the mean and median expression in linear scale separately for cells within the target cluster and for the remaining cells. To ensure numerical stability, a small constant 10^−9^ is added before applying the 𝑙𝑜𝑔_2_ transformation. The 𝑙𝑜𝑔_2_𝐹𝐶 values are then obtained by subtracting the 𝑙𝑜𝑔_2_transformed mean or median expression of the remaining cells from the target cluster. Finally, the results are rounded to three decimal places and stored separately for mean-based and median-based fold-change calculations.
- Area under the ROC curve (AUC). An AUC value of 1 means the gene can perfectly distinguish between the target cluster and other cells, while an AUC of 0 indicates perfect classification in the opposite direction. A value of 0.5 suggests no predictive ability to differentiate between the groups.
- Predictive power score (Power), computed using |𝐴𝑈𝐶 − 0.5| × 2, as implemented in the Seurat package [27]. An AUC of 0.5 means the model has no predictive power, so the difference between AUC and 0.5 reflects the strength of the predictive power. If AUC = 0.5, the result is 0, meaning no predictive power. If AUC = 1, the result is 0.5, indicating perfect prediction. The multiplication by 2 ensures that the final value is in the range of 0 to 1 for ease of interpretability.

### Differential expression analysis

We applied the same differential expression analysis (DEA) methods in the DEA module as in the clustering analysis module for identification of marker genes, with the distinction that the former is not restricted to comparisons based on clustering results. Users can select any combination of available cell annotations for comparison, including the union or intersection of different conditions, allowing for more flexible and customized analyses.

Additionally, we compute two effect size metrics for each gene, the Signal-to-Noise Ratio (SNR), and Cohen’s d. SNR is defined as the ratio of the difference in mean expression between two groups to the sum of their standard deviations. Specifically, for a given gene 𝑔, SNR is calculated as:

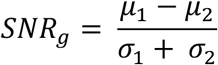

𝜇_1_ and 𝜇_2_ represent the mean 𝑙𝑜𝑔_2_(𝑐𝑜𝑢𝑛𝑡𝑠 + 10^−9^) in the two groups being compared, while 𝜎_1_ and 𝜎_2_ are the corresponding standard deviations. The SNR provides a measure of how clearly a gene’s expression differs between groups relative to the variability within each group, with larger values emphasizing that the difference in expression is more pronounced compared to the noise (variability) in the data. Cohen’s d is defined as the standardized difference in mean expression between two groups, scaled by their pooled standard deviation. Specifically, for a given gene 𝑔, Cohen’s d is calculated as:

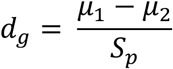

The pooled standard deviation 𝑆_𝑝_ is given by:

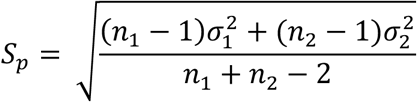

Here, 𝜇_1_ and 𝜇_2_ represent the mean 𝑙𝑜𝑔_2_(𝑐𝑜𝑢𝑛𝑡𝑠 + 10^−9^) in the two groups being compared, 𝜎_1_ and 𝜎_2_ are the corresponding standard deviations, and 𝑛_1_ and 𝑛_2_ denote the number of samples in each group. Cohen’s d provides a standardized effect size, quantifying how many standard deviations apart the two groups are, thereby enabling direct interpretation and comparison across genes, datasets, and studies.

We included both metrics because SNR is less sensitive to unstable variance estimates, avoids division by very small pooled standard deviations (which can inflate Cohen’s d), and behaves more conservatively when one group exhibits high variance, making it a robust metric for feature ranking. In contrast, Cohen’s d provides a standardized measure of effect size, expressed in units of standard deviation, which facilitates interpretation and comparability across studies and datasets.

Our output provides a rich set of metrics, offering multiple ways to assess whether a gene is biologically meaningfully up- or downregulated. We also include pairwise visualizations (e.g., volcano-like plots) to facilitate comparison across metrics. However, interpreting and integrating these metrics manually can be non-trivial. To address this, we define an empirically-inspired weighted-ranking score that integrates significance, magnitude, and effect size into a single value for gene prioritization. Specifically, we combine scaled Cohen’s d, scaled 𝑙𝑜𝑔_2_𝐹𝐶(𝑚𝑒𝑎𝑛), and scaled −log10 adjusted p-values capped at a maximum value of 10, to limit the influence of p-value inflation commonly observed in scRNA-seq data. In addition, we incorporate a noise penalty: for each gene, expression values are subset to the background group, and the maximum expression across those samples is computed, scaled, and subtracted from the final score. The weighting scheme assigns a weight of 2 to Cohen’s d, 0.5 to the p-value component, 1 to 𝑙𝑜𝑔_2_𝐹𝐶(𝑚𝑒𝑎𝑛) and 1.5 to the noise penalty. The aim is to provide a unified metric that captures expression magnitude, statistical significance, and consistency across replicates relative to the contrasting condition. This ranking facilitates the identification of top candidate genes, which can then be further inspected using visualization tools (e.g., violin-plots in the Feature Visualization tab) to assess their suitability as markers of the target condition.

### Functional enrichment analysis

We currently provide two direct methods for scRNA-seq functional enrichment analysis (FEA), gene set scoring and Gene Set Enrichment Analysis (GSEA) [12]. Gene set scores are calculated as the sum of the normalized (log2 median-centered) expression levels of the genes in each user-selected gene set. Users can choose from gene sets included in the MSigDB Collections [30] or define custom gene sets for analysis. A visualization heatmap is available from the markeR package (version 1.1.2) [32] to compare gene sets using two metrics: the Jaccard Index (overlap proportion) and the log odds ratio from Fisher’s exact test (association strength). This feature is particularly useful for comparing user-defined gene sets with reference collections (e.g., MSigDB). GSEA is performed using the fgsea function from the fgsea package [31] (version 1.20.0). Genes are ranked based on a user-selected DEA metric, such as 𝑙𝑜𝑔_2_𝐹𝐶(𝑚𝑒𝑎𝑛) as a measure of magnitude, adjusted p-value for assessing statistical significance, AUC for identifying genes highly specific to each group, and SNR to account for effect size. In addition, we enable users to rank genes according to the predictive power score and absolute value (i.e., module) of the SNR and 𝑙𝑜𝑔_2_𝐹𝐶(𝑚𝑒𝑎𝑛). By focusing on the strength of the signal, regardless of direction, this approach ensures that both upregulated and downregulated genes contribute equally to the detection of biologically meaningful patterns. This is particularly important for identifying changes in pathways where activation or repression involves coordinated expression shifts in both directions. In addition, a dedicated tab is provided for methods that are better suited to bulk data. Within this tab, users can aggregate their dataset into pseudobulk samples (by summation), selecting an appropriate grouping variable and condition. They can then apply score-based approaches implemented in the markeR package [32], including log_2_-median scoring [32], ssGSEA [32], and ranking-based scores [32], a visualize the results using violin-plots of the resulting scores. For each gene set, Cohen’s f is also calculated to quantify the magnitude of differences in scores across groups, providing an effect size measure of how strongly each gene set distinguishes between conditions [32]. Furthermore, the tab also enables evaluation of the discriminatory performance of gene set scores through ROC curve analysis and the corresponding AUC metrics [32].

## Findings

### Key features and functionalities

Extended functionality and diversity of analysis tools are necessary to accommodate the rapidly evolving scRNA-seq research field. scStudio has a **modular structure** that allows straightforward integration of these tools. Each module is dedicated to a specific aspect of scRNA-seq data analysis, from data retrieval and preprocessing to clustering, visualization, and other downstream analyses (Figure 1), namely:

- **Automated retrieval of datasets stored in GEO,** assuming that datasets follow minimal format compliance rules (Figure 2).
- **User count data and metadata** can be uploaded in a wide variety of file formats, including typical .txt, .csv and .tsv formats, 10X Genomics CellRanger count matrix .h5 file and Market Exchange Format (MEX) files (barcodes.tsv.gz, features.tsv.gz and matrix.mtx.gz), as well as Seurat and SingleCellExperiment .rds objects with automatic retrieval of cell-level metadata when available (Figure 2).
- **Integration of multiple datasets**, including datasets stored in different file formats.
- **Flexible data selection, annotation, and subsetting**: scStudio offers robust tools for flexible data handling, allowing users to easily select, annotate, and subset their dataset. This capability enables researchers to tailor their analyses according to specific criteria or experimental needs, enhancing the resolution of their analysis.
- **Comprehensive suite of QC methods**, including flexible parameter selection for usual QC metrics aided with appropriate visualization and tools for identification and removal [20,33] (Figure 3).
- **Library size normalization and batch effect correction** [13,33], with appropriate methods for visual inspection of raw and processed libraries, lacking in most applications (Figure 4).
- **Feature selection and dimensionality reduction methods** with hyperparameter tuning, namely PCA [26], t-SNE [14] and UMAP [15] (Figure 5).
- **Clustering analysis** [28] employing a range of resolutions and a cluster tree [11] to examine how the relationships between cells within each cluster evolve with increasing resolution/ number of clusters (Figure 6).
- **Flexible differential expression analysis** (DEA) to obtain gene signatures for user-defined groups of cells representing cell types, subtypes or states (Figure 7).
- **Comprehensive gene ranking metrics,** complemented by a weighted rank-sum score that integrates significance, magnitude, and effect size for gene prioritization (Figure 7).
- **Functional enrichment analysis** to score user-defined and MSigDB [12,30] gene sets, obtain up- and down-regulated pathways in rankings of genes obtained from DEA of groups of interest, and pseudobulk-based approaches to expand the range of analytical approaches (Figure 8).
- **Extensive visualization methods**, including the distribution of gene expression levels (e.g., violin-plots), gene set scores, average expression in groups of interest (dot-plots), amongst others (Figure 9).
- **History tracking**: for each analysis method applied within scStudio, the system maintains a detailed report documenting the parameters used. This feature ensures that all steps of the analysis are recorded, allowing users to review and track the settings and choices made during the process.
- **Efficient computation management,** by handling time-consuming tasks, such as DEA with large cell groups, through a job submission system. This system processes computations in the server’s background, allowing users to continue exploring and interacting with their dataset without interruption. This feature ensures that extensive analyses do not disrupt the user experience and optimizes workflow efficiency.
- **Session data storage and export**: scStudio enables storage of active sessions, with each analysis step saved as individual .rds files. Users can easily download their entire session, preserving the results, settings and different versions of their analyses.
- **Collaboration and data sharing**: the app supports the publication of data analysis pipelines and facilitates the sharing of private sessions with collaborators. This feature allows researchers to share their work and methodologies transparently, fostering collaboration and enhancing reproducibility within the scientific community.

**Figure 1:**
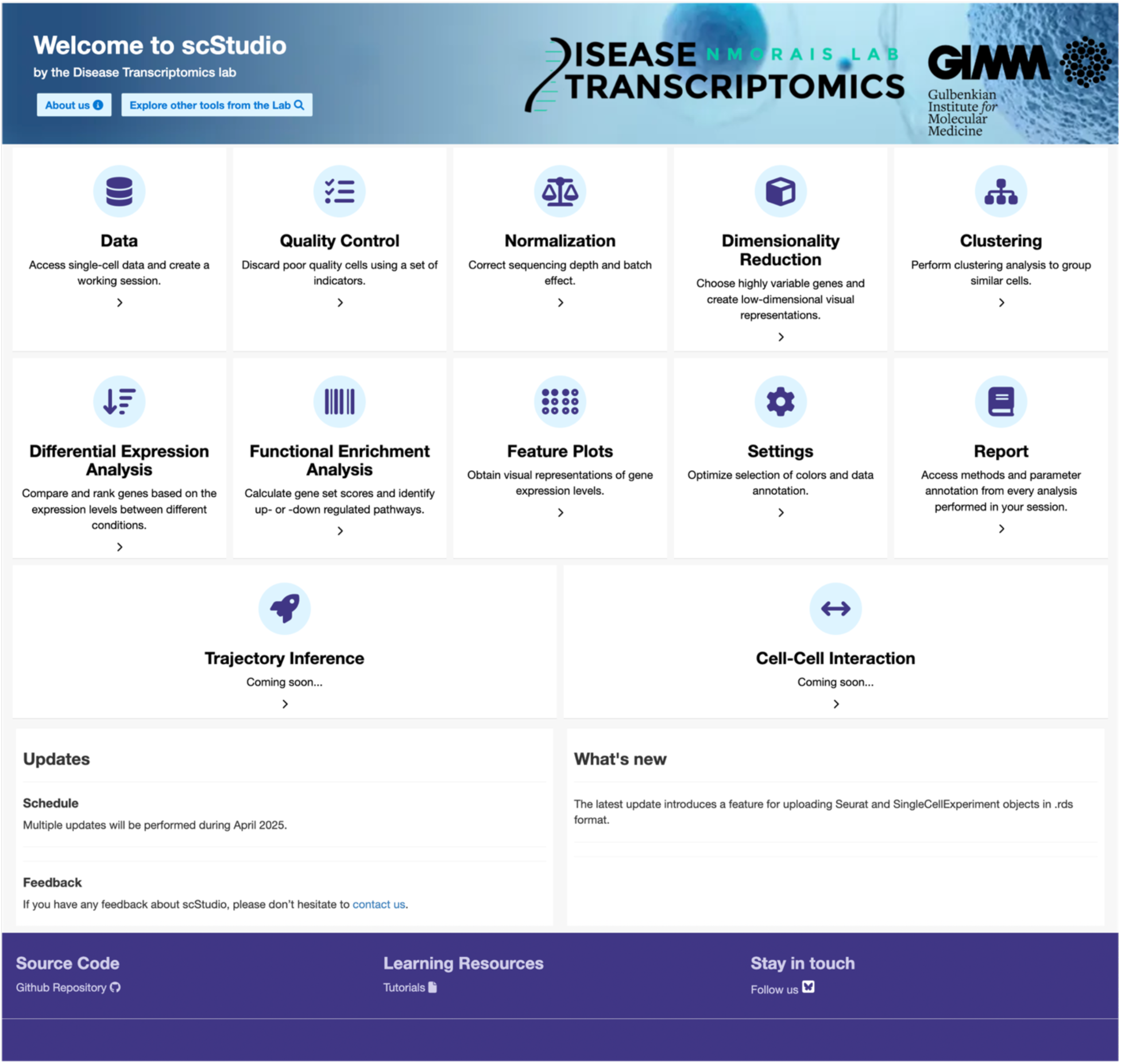
**Entry page of scStudio**, showcasing the available applications for each step of analysis. Users can select tools for data uploading, quality control, normalization, visualization, and downstream analysis. Each app is designed for ease of use, ensuring accessibility for users with varying expertise, while o>ering flexibility to handle complex single-cell datasets.

**Figure 2:**
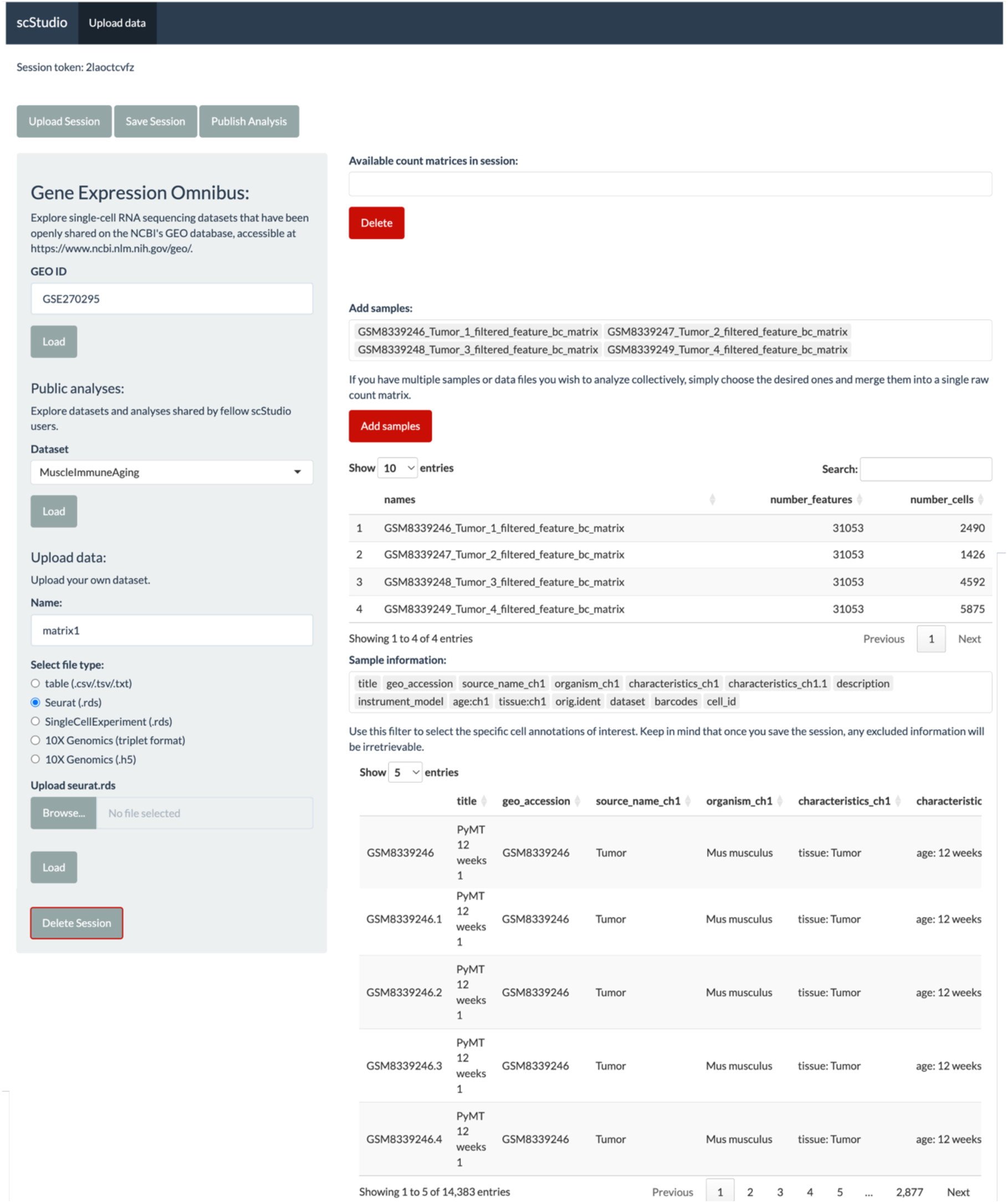
Data Upload Interface. This panel allows users to automatically retrieve data from GEO using unique dataset identifiers. In the example shown, entering ID GSE270295 [47] fetches the associated dataset along with its metadata, streamlining the initial steps of the analysis workflow. It also supports upload of own data in various formats, as well as loading analyses made publicly available by other users.

**Figure 3:**
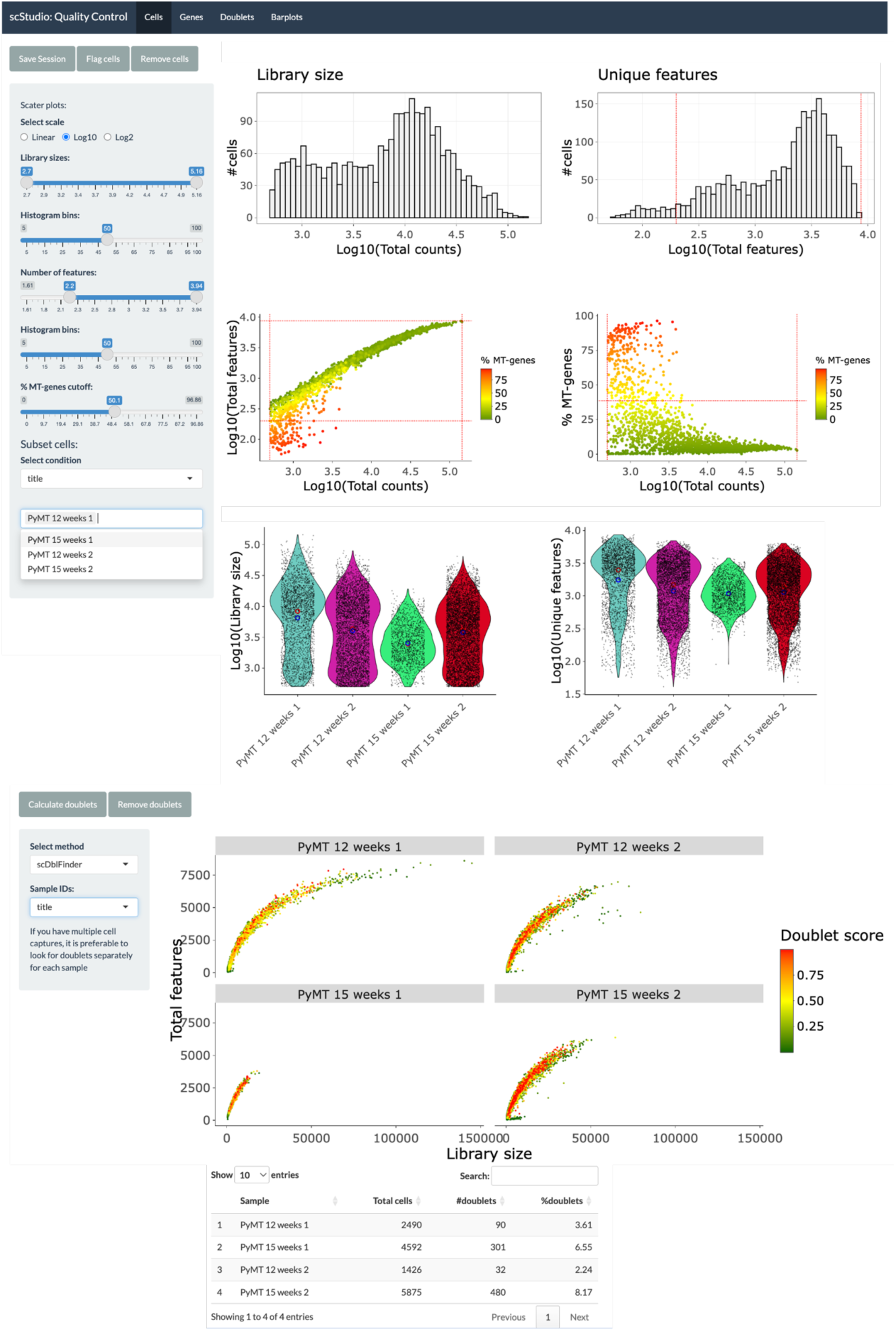
Quality Control Interface. This panel provides examples of comprehensive tools for assessing data quality available at scStudio. Histograms, scatter plots, and violin plots allow users to identify low-quality samples by visualizing library size distribution, number of unique genes detected, and percentage of reads aligned to mitochondrial genes (top). The interface also features doublet identification to detect potential cell doublets (bottom). Users can apply specific quality control measures to individual samples and either flag cells for further inspection or remove poor-quality cells, ensuring robust and accurate data preprocessing tailored to the needs of their study. The example shown uses dataset GSE270295 [47].

**Figure 4:**
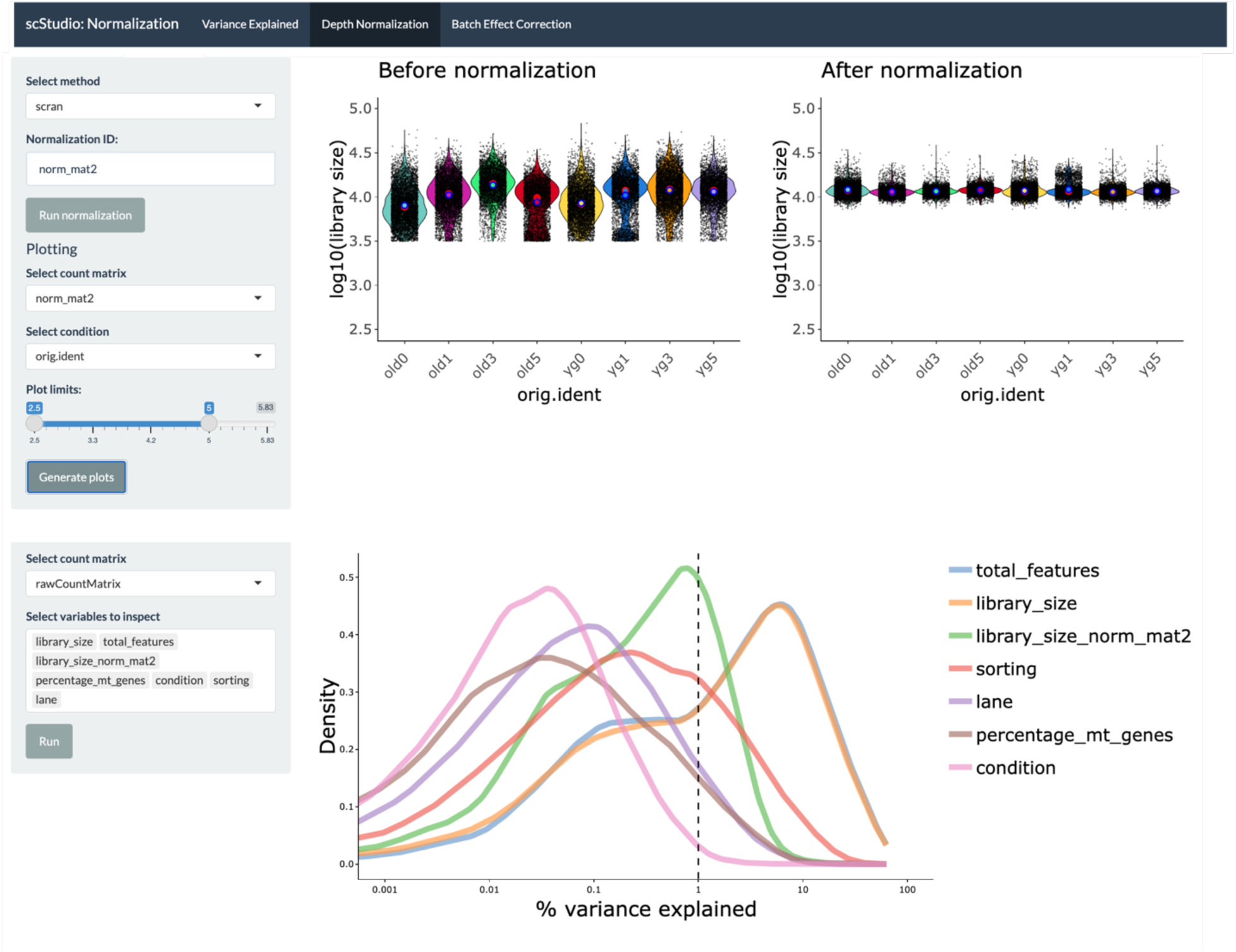
Normalization Interface. The top section displays violin plots comparing the distribution of library sizes across samples before and after normalization, illustrating how the procedure adjusts the data for downstream analysis. The bottom section features a density plot that illustrates the percentage of variance explained by various variables across all genes, including library size before normalization (library_size) and after normalization (library_size_norm_mat2). The example shown uses dataset GSE279741 [48].

**Figure 5:**
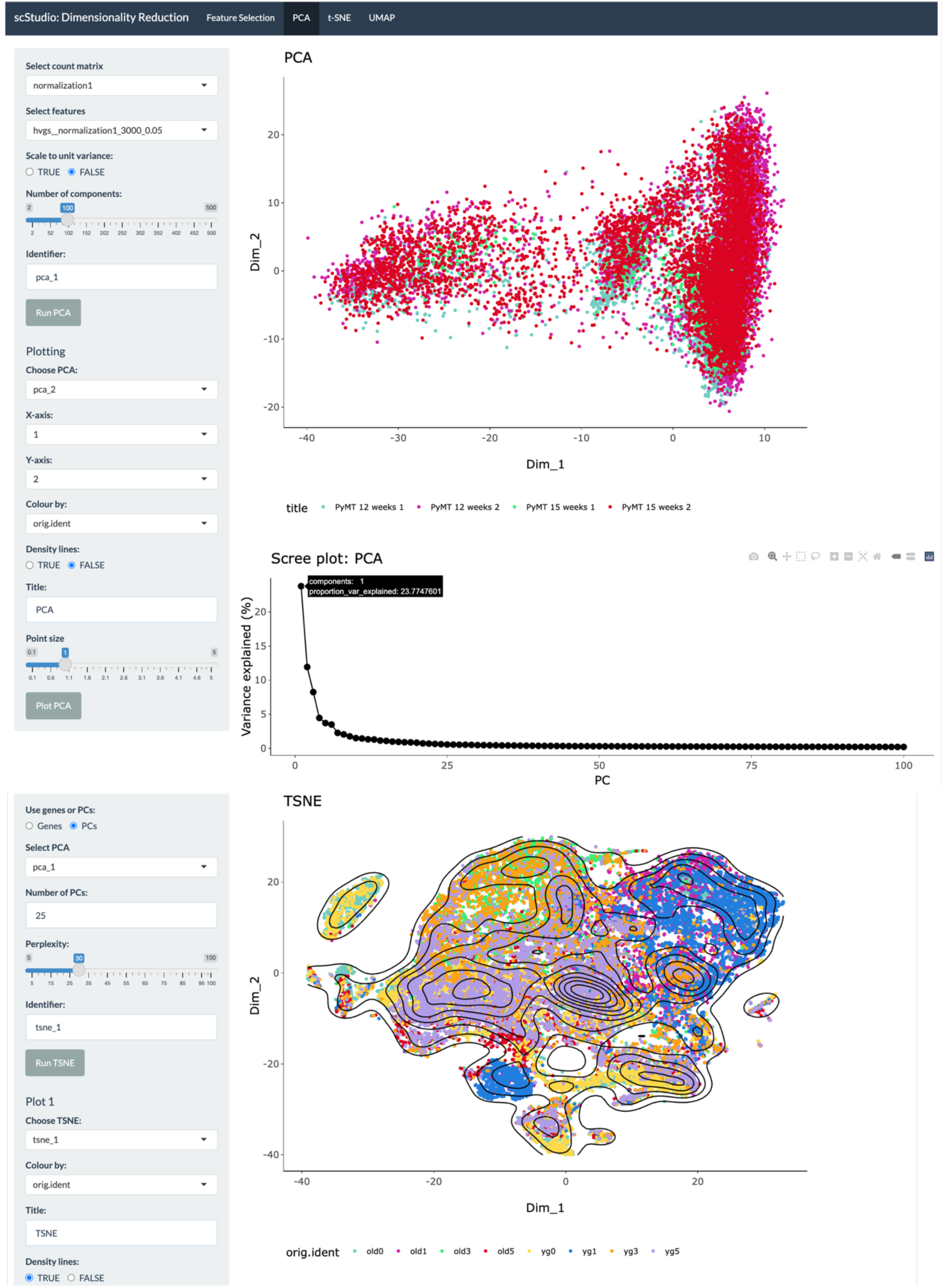
Dimensionality Reduction Interface. The top section features a PCA plot illustrating the major axes of variance in the dataset. A scree plot is included to depict the percentage of variance explained by each principal component, helping users to select the optimal number of components for downstream analysis. The bottom section showcases a t-SNE plot with density lines overlaid, indicating regions with a higher concentration of cells. This visualization is particularly useful for large datasets, where individual cells may overlap, allowing for a clearer interpretation of densely populated areas. The example shown uses dataset GSE279741 [48].

**Figure 6:**
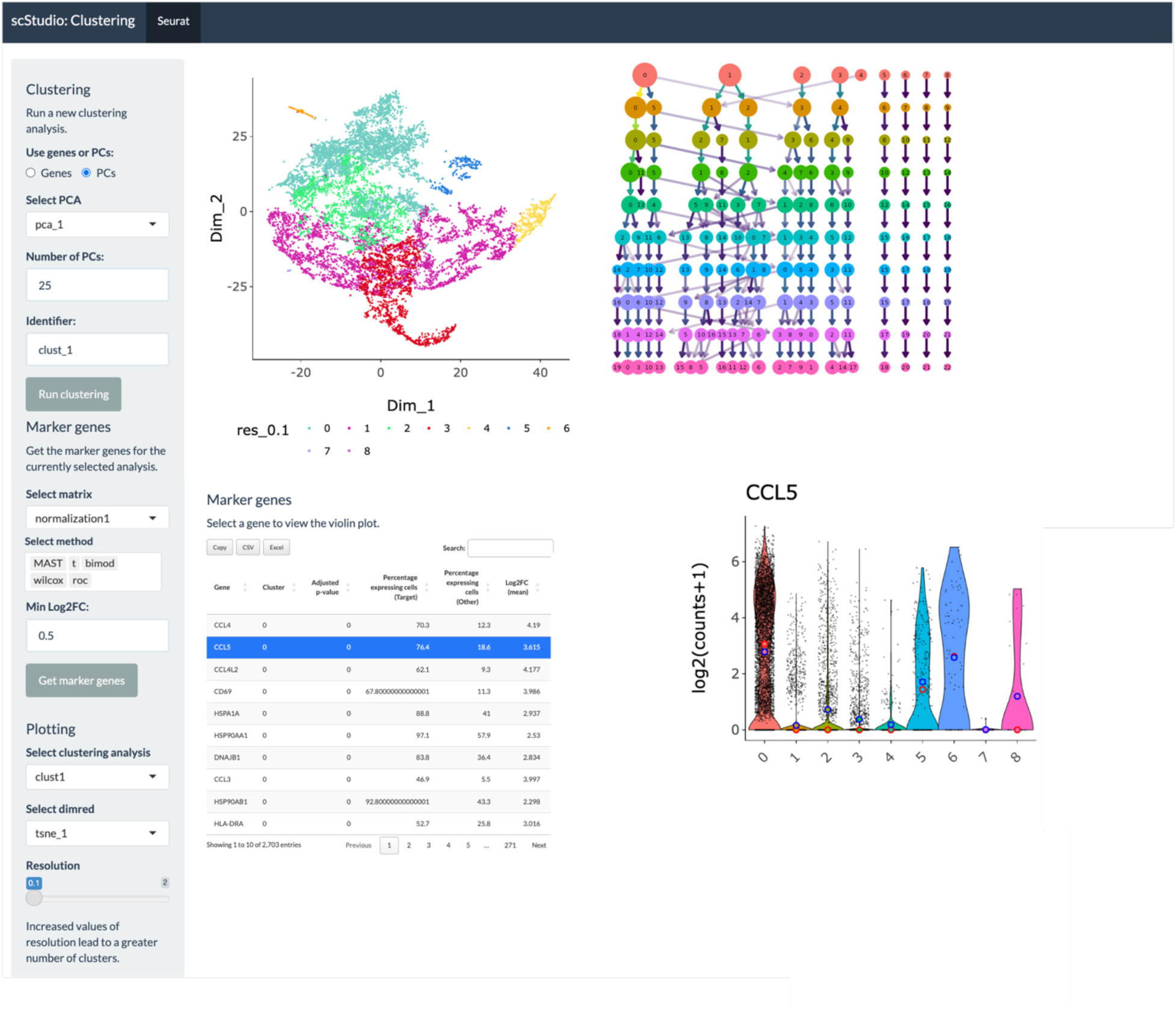
Clustering Interface. This interface includes a user-selected dimensionality reduction plot to display cells colored by their assigned clusters. The cluster tree [11] visually represents hierarchical relationships between clusters, highlighting similarities and potential cell lineages. The accompanying table lists marker genes for each cluster, and users can click on any gene to view its expression pattern as a violin plot. The interface also features a dynamic slider for adjusting clustering resolution, allowing users to explore di>erent levels of granularity. In addition, the user can save multiple clustering results generated with di>erent input parameters, allowing for systematic comparison and optimization of clustering performance. The example shown uses dataset GSE181878 [49].

**Figure 7:**
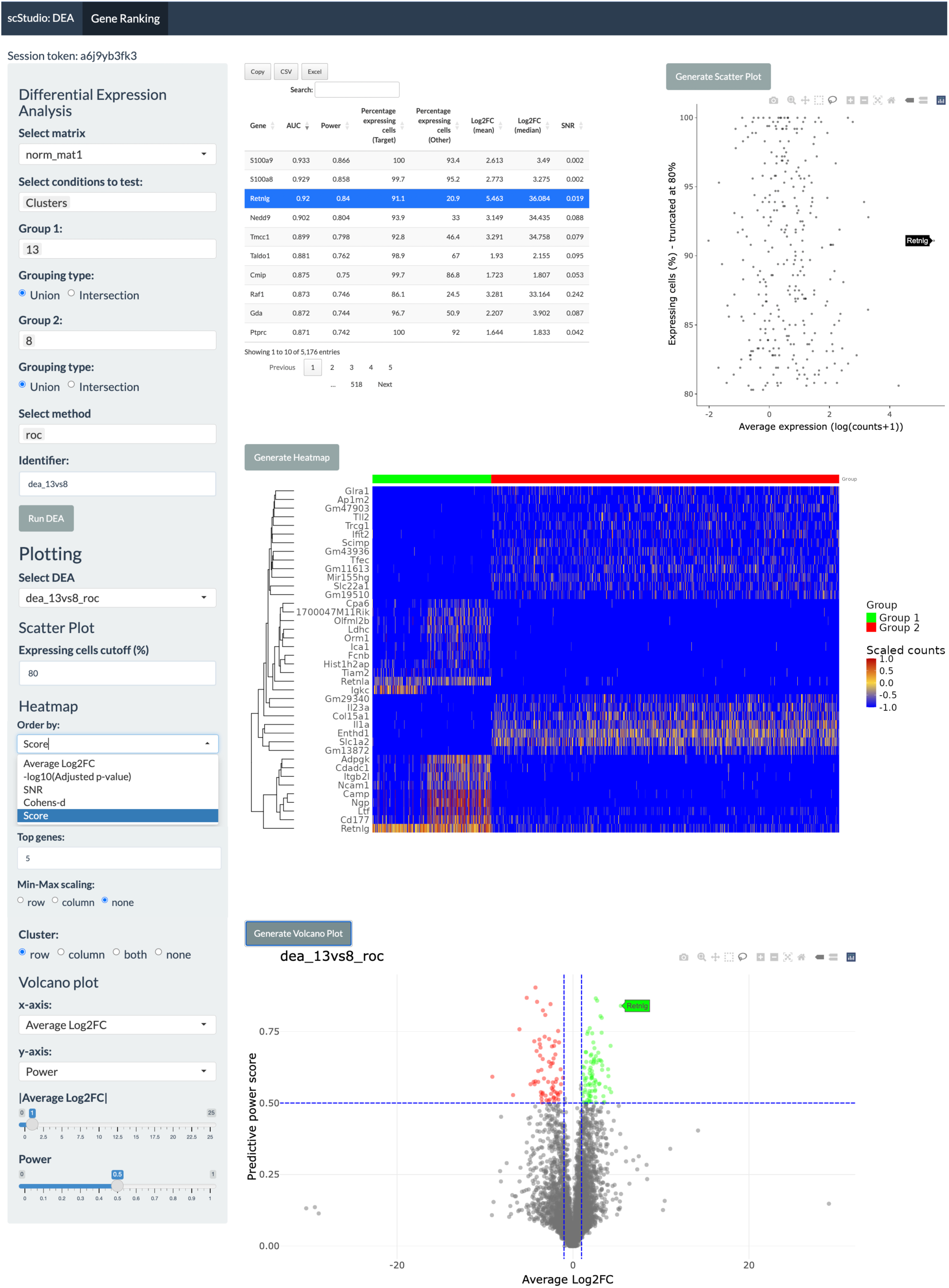
Differential Expression Analysis Interface. This interface provides users with flexible tools to compare user-defined groups of cells, whether based on existing metadata variables or newly created groups defined through direct selection on dimensionality reduction plots. Groups can be refined through unions or intersections of criteria, allowing for nuanced comparisons. The interface features a volcano plot and a scatter plot highlighting genes with high expression levels across a large percentage of cells within the group of interest. An expression heatmap displays selected genes, with customizable options for clustering and scaling. Additionally, a downloadable table of marker genes provides detailed information on gene ranking, aiding in the interpretation of biological significance. The example shown uses dataset GSE279741 [48].

**Figure 8:**
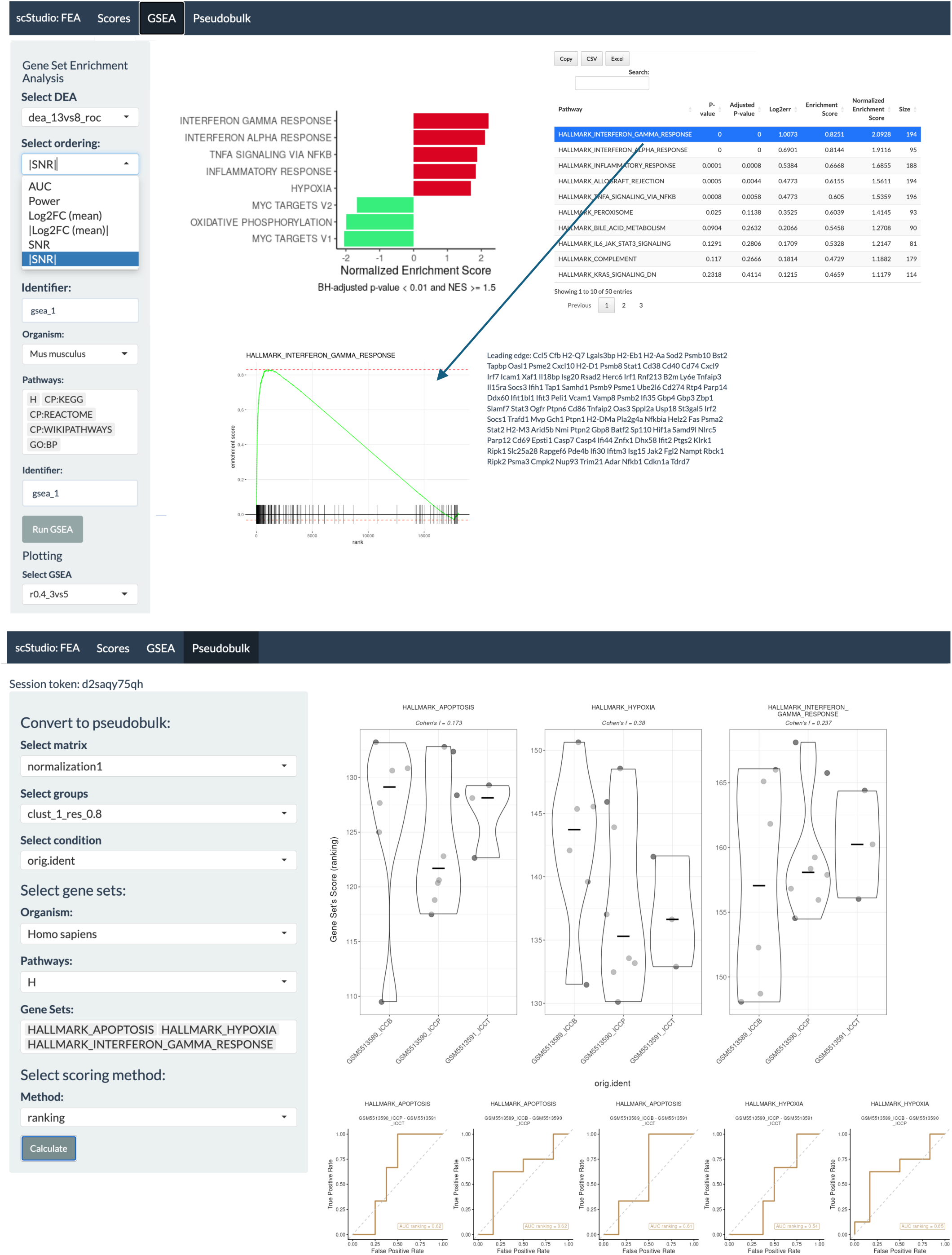
Functional Enrichment Analysis Interface. Users can explore biological pathways through three main options: calculating gene set scores, running GSEA [12] using MSigDB [30] and custom gene sets or converting the dataset into pseudobulk profiles and using the markeR package [32] to evaluate gene sets as phenotypic markers in transcriptomic data. The upper panel includes a bar plot displaying the top up- and down-regulated pathways based on user-defined criteria. Additionally, a ranked table presents pathways with key metrics such as adjusted p-values and Normalized Enrichment Scores (NES). By selecting a pathway from the table, users can view the leading-edge genes contributing to the enrichment, facilitating deeper insights into the biological processes relevant to their data. The lower panel provides a Marker Benchmarking Mode [32], designed to assess the performance of gene sets as markers of a selected metadata variable (e.g., disease state or cellular condition), and to compare results across scoring and enrichment methods. Users can define a grouping variable of interest to aggregate single-cell profiles prior to analysis. The interface includes violin plots displaying gene set scores for the selected condition, computed using log_2_-median expression, ranking-based approaches, and single-sample gene set enrichment analysis (ssGSEA). In addition, ROC curves are generated across contrasts. The example shown uses dataset GSE181878 [49].

**Figure 9:**
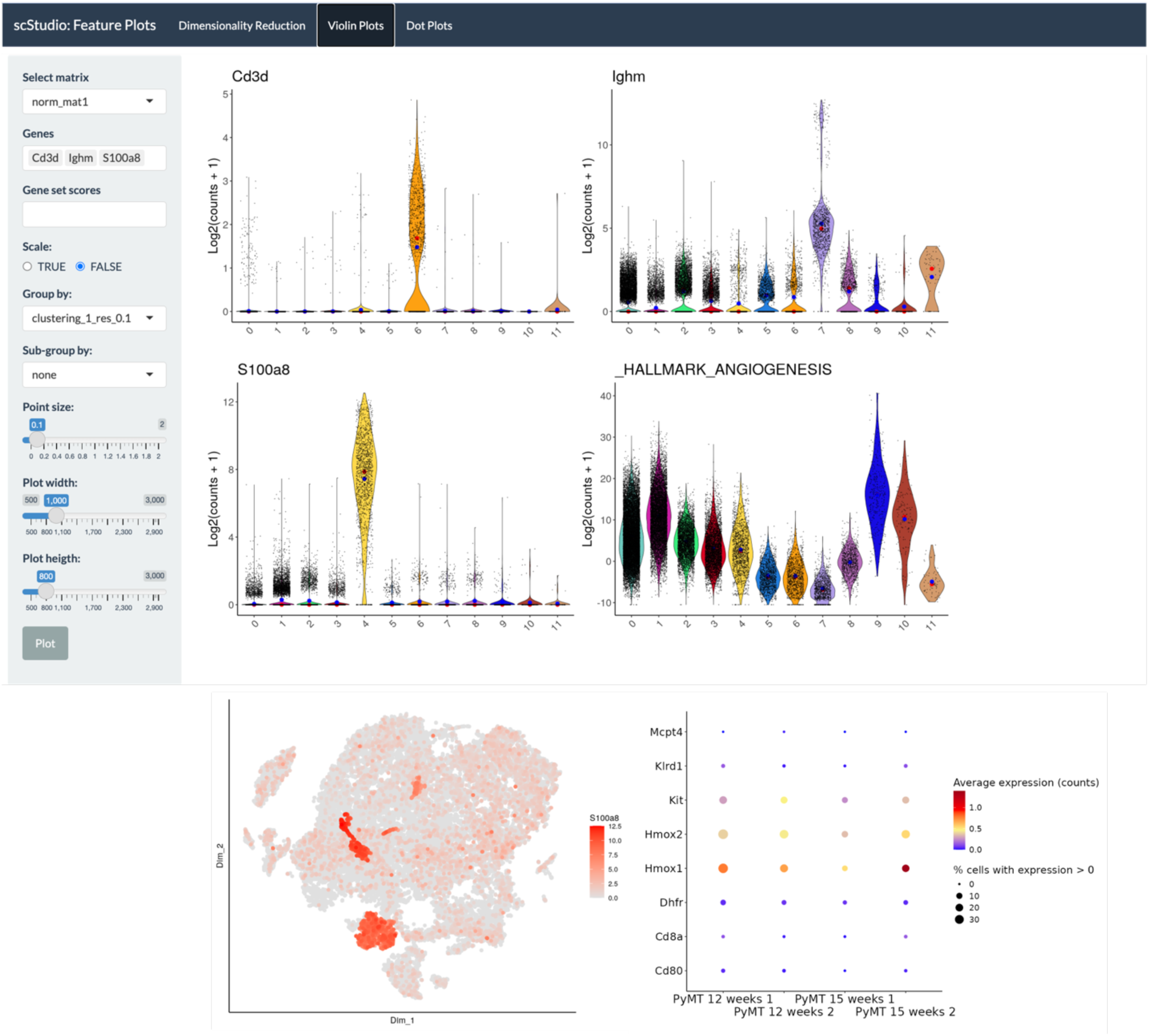
Visualization options in scStudio. scStudio enables multiple ways to explore gene expression data. Violin plots of user-selected marker genes are shown in a grid format, where users can segregate the data based on chosen variables (top). Dimensionality reduction plots, such as t-SNE or UMAP, can be colored by gene expression levels to identify spatial patterns of gene activity (bottom-left). Additionally, a dot plot is available for comparing gene expression averages across groups (bottom-right). These visualization options are also available for gene set scores, allowing for comprehensive exploration and presentation of the data. The example shown uses dataset GSE279741 [48].

### Benchmarking of scRNA-seq data analysis tools

We reviewed several applications projected to facilitate scRNA-seq data visualization and analysis for users without a computational background (Supp. Table 1). For the benchmarking analysis, we selected applications based on the following criteria: the tool must provide a user-friendly **web browser-based interface**, enabling researchers to conduct **comprehensive analyses** without the need for local installations or extensive technical setups; the platform should be **freely accessible** to the public, ensuring that researchers from various backgrounds and institutions can use it without restrictions or licensing requirements; the software should have **open-source code**, promoting transparency, collaboration, and continuous improvement by the community (Supp. Table 1). Three applications stood out as the most complete (Supp. Table 2):

- **ezSingleCell** [35], developed to allow easy integration of scRNA-seq, spatial omics, multiomics, and scATAC-seq data for researchers without computational expertise. For this benchmarking analysis, we focused specifically on evaluating the scRNA-seq module.
- **Automated Single-cell Analysis Portal (ASAP)** [36] developed to facilitate the comprehensive and scalable analysis of scRNA-seq data, making it accessible for researchers without a computational background.
- **ICARUS (version 3)** [37], featuring a tutorial-like interface that enables users without programming knowledge to perform scRNA-seq data analysis.

Among the three applications, ASAP stands out as the only one that provides access to public databases. However, its current functionality is limited to cloning projects from existing datasets in the Human Cell Atlas [38] and the Fly Cell Atlas [39], which comprise 56 datasets, along with 93 projects shared by users. In contrast, scStudio facilitates automatic access to GEO [8], including datasets from these databases, while also expanding the scope to most publicly available single-cell datasets (Figure 2).

Although users can manually upload count data, ezSingleCell and ICARUS do not support the 10X Genomics CellRanger output files in MEX, commonly stored in GEO. Furthermore, their support for other file formats is limited; for instance, ICARUS requires specific sample annotation when uploading multi-sample datasets, which may require *a priori* data manipulation. Additionally, neither ezSingleCell nor ICARUS allow the upload of metadata, such as tables for cell-level variable annotations.

scStudio supports uploading and integration of multiple datasets, independently of file format, a feature not available in ASAP and limited to two datasets in ICARUS. We also found that none of the three applications allow in-depth quality control to identify and filter poor quality cells or discriminate poor quality samples. For example, none of the applications supports segregation of violin plots to visualize the distribution of library sizes, the number of unique genes, and the percentage of reads aligned to mitochondrial genes plots <u>by</u> sample or other relevant categorical variable, such as batch (Figure 3). Additionally, the tools lack the ability to plot data on a log scale, which is crucial for identifying deviations from the expected log-normal distribution of quality metrics. This feature is especially important for improving the resolution of low-end metrics, where, aside from mitochondrial genes, poor-quality cells are typically concentrated (Figure 3). Moreover, ezSingleCell does not allow filtering cells based on library sizes, which is a critical step in quality control.

Neither ASAP nor ezSingleCell include tools for doublet identification and removal, which is an essential step in the quality control of scRNA-seq data generated using droplet protocols [29]. Furthermore, none of the applications offers visualization methods to assess library size normalization and batch effect correction results. For example, they do not provide violin plots that illustrate the distribution of library sizes before and after normalization, or the variance explained by library sizes and other potential sources of batch effect in the data (Figure 4). In addition, ezSingleCell does not offer an option for intra-dataset batch effect correction, such as regressing out confounding variables e.g., library-preparation batches. For downstream analysis, it is crucial that the process remains transparent rather than operating as a “black box”, while also being flexible enough to accommodate the visualization of parallel analyses. For instance, users should be able to choose between unnormalized, normalized, and batch-effect corrected count matrices, as well as specify the number of genes (e.g., HVGs) and the scaling/centering of the matrix when applicable. Additionally, users should have access to visualization tools that aid in parameter selection, such as scree plots for determining the appropriate number of principal components (PCs) for dimensionality reduction techniques and cell clustering (Figure 5), or cluster trees [11] (Figure 6) to assist in selecting the optimal clustering resolution. These are features that are provided in scStudio but fail to be available in some of the apps that we tested. For example, ASAP does not provide a scree plot nor a cluster tree. The latter feature is also unavailable in ICARUS.

We also noted that among the three applications, the scaling and centering of the count matrix for PCA are handled differently: ICARUS automates this process, ASAP does not allow centering without scaling and ezSingleCell does not provide information regarding this step. Furthermore, users are restricted to performing dimensionality reduction methods and clustering analyses using principal components (PCs) rather than the count matrix itself. In contrast to scStudio, that automatically offers a window of clustering resolutions for dynamic inspection (Figure 6), the three benchmarked apps also provide only one user-selected resolution at a time. Additionally, in ICARUS, each new resolution result overwrites the previous one.

Another key feature is the availability of a wide variety of flexible plots for visualizing gene expression. For example, scStudio offers violin plots that can be segmented by any categorical variable in the dataset’s metadata, as well as dot plots that show both average expression and the percentage of cells expressing a user-selected gene (Figure 9). Users can also visualize scores based on gene sets they manually annotate or that are automatically retrieved from the MSigDB collections [12, 30] (Figure 9). Aside from scStudio, only ICARUS offers similar capabilities.

While all applications enable users to perform FEA using gene ranks obtained through DEA, ezSingleCell only allows users to perform this analysis using the groups from the automatic cell type identification step. scStudio not only enables flexible analysis with user-selected groups but also provides a table with analysis statistics, such as adjusted p-values and normalized enrichment scores (NES) [12]. It also allows users to inspect leading edge genes, which are the genes contributing to the enrichment score of the up- and down-regulated pathways [12] (Figure 8).

Finally, only scStudio and ASAP allow users to run parallel pipelines. However, ASAP requires users to log in with an account ID and password to store sessions. In contrast, scStudio assigns a unique token to each analysis, allowing users to easily save, clone and share analyses directly with collaborators, without requiring them to have an account. Finally, both scStudio and ASAP allow users to either store the analysis on a dedicated server or download it to their personal computers, while ICARUS allows users to download the analysis and ezSingleCell does not have an option for users to save their working session in any format, which hinders progress and data sharing. Other applications included in our list (Supp. Table 1) are primarily data visualization tools and do not allow to perform in-depth quality control and downstream analysis of single-cell data. To our knowledge, only clustifyr [40] allows automated access to GEO single-cell datasets. However, the purpose of this application is focused on rapid automatic cell type annotation of scRNA-seq data using a library of cell type references [40], hence lacking tools for a comprehensive data analysis. In comparing these tools, we found that while each platform has its strengths, scStudio offers a comprehensive suite of tools that make it particularly well-suited for flexible, customizable single-cell RNA-seq analyses, supporting both novice and experienced users.

## Discussion

### What defines an effective scRNA-seq analysis tool?

Latest developments in molecular profiling technologies resulted in the creation of large scRNA-seq datasets, many of those not fully explored yet, but accessible to the scientific community via public data repositories. Previous attempts have been made to create user-friendly tools for analyzing single-cell data without requiring computational expertise (Supp. Table 1). However, many existing tools lack comprehensive visualization methods for deep analysis and offer limited flexibility in data handling. Additionally, these tools typically require users to provide data in specific file formats, such as .txt count tables. This requirement could pose a challenge for scientists lacking programming skills, especially considering that the main database for publicly shared single-cell data i.e., GEO [8], does not enforce a particular format for processed and supplementary (e.g., metadata) data submission. With scStudio we aim to contribute towards bridging the conceptual and knowledge gaps between wet- and dry-lab researchers, by making bioinformatics tools and database access freely available and easily usable by the entire scientific community. We focus on areas that are vital to that goal, namely, creating a virtual infrastructure to support the active sharing of bioinformatic tools developed for scRNA-seq data analysis to the broad scientific community. We also provide automated access to GEO [8], the largest public single-cell data repository, with the goal of facilitating the reanalysis of published datasets. Automated retrieval not only saves time but also allows for standardized and reproducible data acquisition processes. The retrieved data can then be processed and prepared for analysis using the built-in tools provided by scStudio, streamlining the workflow for users and minimizing the need for external software.

### Which data formats are typically published on GEO, and which ones are supported by scStudio?

Currently, scStudio allows automated retrieval of datasets stored in GEO [8] using the most common data file formats. These include the following: CellRanger output MEX files, .h5 files and standard .txt, .csv and .tsv count table files. During the development of scStudio, we encountered challenges stemming from the lack of strict rules for data sharing on GEO. Current guidelines [41] allow supplementary files to be included without specification of the format. This can lead to cases where data is shared in suboptimal formats such as Excel files with lengthy text headers (<u>GSE103840</u>). These formats can present difficulties in programmatically uploading data to standard data analysis interfaces like R and Python. Moreover, some studies store multiple modalities of single-cell data within a single GEO entry. For example, it is not uncommon to find spatial transcriptomics (<u>GSE243275</u>), CITE-seq (<u>GSE201809</u>) or ChIP-seq (<u>GSE179080</u>) data stored alongside scRNA-seq data, among others. Furthermore, some entries combine scRNA-seq experiments conducted with different protocols, complicating sample annotation and making it more challenging to apply necessary batch effect correction methods (<u>GSE139627</u>; <u>GSE166548</u>; <u>GSE163108</u>; <u>GSE131309</u>). Additionally, there seems to be a lack of oversight regarding the review of stored data files. We came across a few instances where individual files did not have any proper labelling/sample identification (<u>GSE175520</u>; <u>GSE223373</u>), highlighting the need for more thorough quality control measures.

Historically, GEO was developed primarily to store and provide access to microarray data [8]. It has since expanded to accommodate various types of high-throughput sequencing data, including bulk and scRNA-seq, among others. Since GEO was originally designed for data where samples typically represent an entire experiment or condition, metadata is commonly stored at the sample level rather than the more granular cell level. This historical design can lead to challenges when working with single-cell data, where metadata at the cell level is often crucial for comprehensive analysis and interpretation. In response, scStudio automatically collects available sample-level data annotation and converts it at the cell level, facilitating comparisons between groups of interest and other relevant variables (such as technical factors stemming from various processing batches). However, this storage approach typically excludes crucial downstream analysis results, such as clustering analysis and cell type annotations. Puntambekar and colleagues [10] estimated that less than 25% of studies provide useful metadata at the cell level, including identified cell types. This poses significant challenges for reproducibility efforts, knowledge transfer and leveraging the wealth of public data. Füllgrabe *et al*. [42] proposed minSCe - Minimum Information about a Single-Cell Experiment, a framework to provide a minimum set of metadata categories and checklist designed to ensure consistency in reporting scRNA-seq data. This includes critical information that describes a scRNA-seq assay in sufficient detail to facilitate analysis and replication [42]. A key feature is the unique cell identifier. Since cell barcodes can potentially be duplicated across datasets and samples, having a distinct cell identifier is essential for tracing any procedure back to the original cell. scStudio addresses this by generating random *de novo* cell IDs as soon as data are uploaded to the platform, in addition to storing the original barcodes, sample and dataset annotation. Another key feature is the “inferred cell type” based on gene expression patterns. As this classification is determined post-analysis, the framework emphasizes the importance of documenting reproducible analysis steps to enhance transparency and support scientific scrutiny [42]. While previous studies have raised awareness of this issue [10,42], there has been a notable lack of progress in enhancing GEO’s single-cell data storage in recent years. There have been other initiatives aimed at creating efficient single-cell data databases, such as the Single Cell Portal [43] and PanglaoDB [44], although the latter is no longer actively updated. scStudio addresses this by enabling users to upload a high diversity of files directly downloadable from these databases (e.g., Seurat .rds), including metadata files.

### Challenges of automation and the importance of providing comprehensive tutorials

The analysis of a scRNA-seq data should be personalized to each individual dataset given their diversity in terms of quantitative (number of samples and number of cells) and qualitative characteristics (condition, sample type, protocol for tissue dissociation and library preparation, expected cell type diversity, amongst others). Every step of the analysis involves selecting appropriate parameters, e.g., cutoff values for quality control, identifying and removing potential batch effect sources and selecting a reasonable number of clusters that accurately reflect groups of cells with a relevant biological function. It is important to revise these decisions through the course of the downstream analysis in an iterative manner. scStudio was developed as a **decision-support tool** that allows users to maintain a record of the parameters selected for each analysis, as well as conducting parallel pipelines with variations of the selected tools and parameters. For example, in scStudio we apply the Bonferroni method to correct p-values for multiple comparisons in differential expression analysis. This is a conservative correction compared to e.g., Benjamini-Hochberg’s false discovery rate, to help reduce the risk of detecting statistically significant but biologically irrelevant differences. Nonetheless, even with this correction the resulting p-values remain very low and could be misinterpreted, since p-values tend to approach zero in very large samples even for negligible effects [45]. Therefore, statistical significance alone should not be overinterpreted and proper evaluation of effect size metrics is essential to avoid misleading conclusions. Finally, methods should not be completely automated and devoid of user intervention. Instead, users should be knowledgeable and informed about each step. Adjusting parameters allows to account for differences and optimize the analysis for each specific dataset. Methods with multiple tunable parameters can be extremely useful but also depend on developers providing clear guidelines on how to set them. To guide users through scStudio’s multiple functionalities and key considerations for scRNA-seq data analysis, we provided high-quality tutorials and training videos on using scStudio and sharing results with the community. These materials were designed not only for **inexperienced** users, but also for **more advanced users**.

## Conclusion

Our work aims to consolidate the fundamental FAIR principles – Findability, Accessibility, Interoperability, and Reusability [46] by enhancing the management of single-cell transcriptomic data. scStudio embodies these principles, enabling all biomedical scientists, including those without programming expertise, to easily retrieve datasets from repositories housing most publicly available single-cell data. Future expansions of scStudio will integrate additional databases, such as the Human Cell Atlas [38]. Additionally, scStudio empowers researchers to conduct transparent and in-depth quality control and exploratory analyses of scRNA-seq data, providing visual tools to guide intelligible decision-making and thereby avoid the use of ‘black-box’ methods. The platform facilitates not only the integration of multiple datasets and user data but also the sharing of analysis pipelines and annotations within the scientific community. A key aspect of scStudio’s success will be its ability to integrate diverse tools and approaches developed across different laboratories, generating a platform for community feedback, interdisciplinary dialogue, and fostering new collaborations.

## Availability of Supporting Source Code and Requirements

Project name: scStudio

Project home page: https://github.com/DiseaseTranscriptomicsLab/scStudio Operating system(s): Platform independent

Programming language: R Other requirements: None

License: GNU General Public License v3.0 RRID: SCR_027618

bio.tools ID: scstudio

## Additional files

Supplementary Table 1.

Supplementary Table 2.

## Abbreviations

ASAP: Automated Single-cell Analysis Portal
AUC: area under the ROC curve
CSV: comma-separated values
DEA: differential expression analysis
FEA: functional enrichment analysis
GEO: Gene Expression Omnibus
GSEA: gene set enrichment analysis
HVGs: highly variable genes
MEX: Market Exchange Format
NES: normalized enrichment score
PCA: principal component analysis
PCs: principal components
QC: quality control
ROC: receiver operating characteristic analysis
scRNA-seq: single-cell RNA sequencing
SNN: shared nearest neighbor
t-SNE: t-distributed stochastic neighbor embedding
TSV: tab-separated values
TXT: plain text
UMAP: uniform manifold approximation and projection
UI: user interface
XLSX: Excel files

## Authors’ contributions

M.B. and N.L.B.M. conceived the project, M.B. developed the tool, benchmarks and wrote the manuscript draft. N.L.B.M. and K.S. discussed the results and revised the manuscript.

N.L.B.M. supervised this work.

